# A Cell-Autonomous Role for the Vitamin B6 Metabolism Gene *PNPO* in *Drosophila* GABAergic Neurons

**DOI:** 10.1101/2025.08.07.669010

**Authors:** Wenqin Fu, Saul Landaverde, Xiaoxi Zhuang, Atulya Iyengar

**Affiliations:** Department of Biology, and Iowa Neuroscience Institute, University of Iowa, Iowa City, IA 52242, USA; Department of Biological Sciences, University of Alabama, Tuscaloosa, AL 35487, USA

**Author notes:** Corresponding author: Atulya Iyengar, Department of Biological Sciences, University of Alabama, Tuscaloosa, AL 35473.

**Keywords:** Pyridoxal, Pyridoxal 5’-Phosphate

## Abstract

In animals, the enzyme pyridox(am)ine 5’-phosphate oxidase (PNPO) is critical for synthesizing the active form of vitamin B6 (VB6), pyridoxal 5’-phosphate (PLP), from inactive vitamers. PLP is a required cofactor for many enzymatic reactions, including the synthesis of GABA and the monoamines. *PNPO* disruption in humans is associated with an array of epilepsy syndromes, while *Drosophila* harboring mutations in the sole *PNPO* ortholog, *sugarlethal* (*sgll*), display spontaneous seizures and short lifespans. These phenotypes are suppressed by PLP supplementation and are exacerbated by restriction of dietary B6 vitamers. In the context of PNPO deficiency, it remains to be resolved what the specific contributions by cellular subpopulations in the nervous system are to neurological phenotypes. We addressed this question in *sgll* mutants by expressing human *PNPO* (*hPNPO*) cDNA in cholinergic, glutamatergic, and GABAergic neurons as well as glia and measuring changes in survival and seizure phenotypes. We found *hPNPO* expression in GABAergic neurons largely restored lifespan and attenuated seizure activity, while glial expression also improved *sgll* phenotypes albeit to a lesser degree. In contrast, *hPNPO* expression in either cholinergic or glutamatergic neurons, accounting for most neurons in the fly brain, did not appreciably alter *sgll* phenotypes. We contrasted these observations with changes in *sgll* mutants induced by feeding GABA receptor modulators. The GABA_B_ agonist SKF-97541 reduced mortality, while GABA or GABA_A_ receptor modulators did not improve survival. Together, our data establish a cell-autonomous role for PNPO in GABAergic neurons to support brain function, especially under VB6-restricted conditions.

## Introduction

The enzymatically active form of vitamin B6 (VB6), pyroxidal 5’ phosphate (PLP), is a necessary cofactor for over a hundred biochemical processes across the tree of life. Plants, bacteria, archaea and fungi can synthesize PLP *de novo* [1]. Animals must instead scavenge VB6 vitamers pyridoxine (PN), pyridoxamine (PM) or pyridoxal (PL), and their 5’ phosphorylated species (PNP, PMP and PLP respectively) from their diet [2, 3]. Two enzymes form a biochemical pathway to salvage these inactive VB6 vitamers into PLP in animals: pyridoxal kinase (PDXK), which phosphorylates the B6 vitamers; and pyridox(am)ine 5’-phosphate oxidase (PNPO), which oxidizes PNP/PMP into PLP [4, 5]. Under physiological conditions in animals, PNPO serves as the rate-limiting step in PLP synthesis.

PLP plays a critical role in neurotransmitter metabolism. Enzymes involved in the synthesis of GABA (glutamate decarboxylase); dopamine, serotonin, epinephrine (aromatic L-amino acid decarboxylase); and histamine (histadine decarboxylase) all require PLP as a co-factor [6]. In humans, mutations in the PNPO enzyme which lead to reduced PLP levels are associated with a variety of neurological disorders including neonatal epileptic encephalopathy and other epilepsy syndromes [7-11]. Consistently, disruption of *PNPO* orthologs in zebrafish (*pnpo*, [12, 13]) and *Drosophila* (*sugarlethal, sgll*, [14]) lead to spontaneous and recurrent seizures. In fly *sgll* mutants, both dietary PLP supplementation or transgenic expression of human wildtype *PNPO* cDNA (*hPNPO*) can attenuate seizure phenotypes [14, 15].

An important question regarding genotype-phenotype relationships in *PNPO* mutants is the nature of PNPO function in specific tissues of the nervous system. Conceivably, interconversion between cell-permeable PL and non-permeable PLP (via PDXK) may be sufficient in certain cell types, while in other cells PNPO-mediated PLP production is necessary for proper function. In flies, pan-neuronal RNAi knock-down of *sgll* transcripts leads to substantially milder seizure phenotypes compared to knock-down across all tissues and further restrictions to neuronal sub-types do not yield obvious phenotypes [14]. These observations suggest PL import and interconversion to PLP partially modulates seizure expression. However, the contributions of disrupted VB6 metabolism in specific cell types to the emergence of seizures and other neurological phenotypes remain to be delineated.

Cell-type specific manipulations of PNPO expression could reveal tissues critical for generating seizure phenotypes associated with *PNPO* mutations. Here, we examined how cell type-specific *hPNPO* expression (via the gal4 – UAS system [16]) modulated seizure phenotypes in the fly *sgll*^*95*^ mutant. Under VB6-deprived conditions, *sgll*^*95*^ flies display spontaneous seizures and die within 3-5 d. We examined the degree to which human *PNPO* cDNA (*hPNPO*) could attenuate survival and seizure phenotypes when expressed in glia, as well as in cholinergic, glutamatergic, and GABAergic neuron sub-populations in *sgll*^*95*^ mutants. Our findings indicate survival and seizure phenotypes are largely attenuated by *hPNPO* expression in GABAergic neurons and to a lesser degree, in glia. In contrast, *hPNPO* expression in cholinergic neurons (the primary excitatory neurons in *Drosophila*) or in glutamatergic neurons did not appreciably modulate these phenotypes.

## Methods

### Fly Stocks

The *sgll*^*95*^ line and the reference line *w*^*1118*^ are described in Chi et al. (2014)[17] and (2019)[14]. The UAS *hPNPO* line expressing human *PNPO* cDNA is described in Chi et al. (2019)[14]. The gal4 driver lines *GAD* (BL #51630), *VGLUT* (BL #26160, also known as *OK371* gal4), *ChAT* (BL # 6793) and *repo* (BL # 7415) were obtained from the Bloomington *Drosophila* Stock Center (BDSC) at Indiana University. To generate lines expressing UAS *hPNPO* under the control of a gal4 driver in the background of *sgll*^*95*^, we first generated lines of the UAS *hPNPO* and gal4 drivers on a *sgll*^*95*^ background via classical fly husbandry (UAS *hPNPO/*CyO ; *sgll*^*95*^ and gal4/CyO ; *sgll*^*95*^). The UAS *hPNPO* and gal4 lines were then crossed with each other to generate flies of interest. Control flies were generated by crossing these lines with the original *sgll*^*95*^ background lines as indicated. Crosses were set-up at room temperature (∼23 °C) under a 12h : 12h light-dark cycle in fly vials containing standardized cornmeal media diets. (The cornmeal-yeast-molasses diet was used at U. Chicago [15], Bloomington fly food recipe was used at U. Alabama.) Following eclosion (< 2 d), flies were collected over CO_2_ anesthesia and reared on VB6-free media consisting of 4% sucrose, 1% agar [17].

### Survival Analysis and Pharmacology

Male flies were collected in 0-2 d range, placed on sucrose-only diet in groups of < 20 flies/vial and survival was checked daily (see Chi et al., 2019 for details). For pharmacological experiments (Figure 3), GABA (Millipore Sigma A2129), Diazepam (Millipore Sigma D-0899), SKF-97541 (Millipore Sigma 5097080001) or Gaboxadol (Millipore Sigma #16355) were mixed into the VB6-free media at the indicated concentrations.

### Electrophysiology

Tethered fly electrophysiology was performed as described in Chi et al., 2019 [14] and Iyengar & Wu, 2014 [18]. Male and female flies reared on VB6-free media as described and aged 3-5 d were observed. Flies were anesthetized on ice and fixed to a tungsten pin with UV glue (Locktite #4311). Following a ∼30 min recovery period, electrolytically sharpened tungsten electrodes were inserted into the top-most dorsal longitudinal muscle (DLM, #45a in [19]) on the left and right sides. A similarly constructed electrode inserted into the dorsal abdomen served as the reference. Intracellular DLM spiking was picked up by an AC amplifier (AM Systems Model 1800) and digitized by a data acquisition card (National Instruments USB 6210) controlled by LabView (version 2022, National Instruments). Spike detection and analysis was done offline using custom-written Matlab scripts (r2022b, Mathworks). Overall DLM firing rate and DLM bursting were determined as described in Lee et al., 2019 [20].

### Statistics

All statistical analyses were performed in Matlab using the Statistics and Machine Learning Toolkit. Details on sample size, statistical tests performed, multiple test corrections, and statistical significance are provided in the figure legends or in text describing each figure.

## Results

### Cell-type specific expression of hPNPO in *sgll* mutants modulates survival on VB6-free media

We have previously shown that *sgll*^*95*^ mutant flies display increased mortality (3-5 d) when reared on a VB6-free diet, and in these mutants, expression of the UAS *hPNPO* construct under the control of the ubiquitous *act5c* gal4 (i.e. UAS *hPNPO / act5c* gal4 ; *sgll*^*95*^) rescued the mortality phenotype [14]. Using a similar strategy, we sought to determine if UAS *hPNPO* expression restricted to cholinergic neurons (via *ChAT* gal4), the primary excitatory neurotransmitter in the fly brain, accounting for ∼ 60% of all synapses; glutamatergic neurons (via *VGLUT* gal4) which often function as motor neurons or inhibitory interneurons; GABAergic neurons (via *GAD* gal4), the primary class of inhibitory interneuron; or glia (via *repo* gal4), accounting for ∼ 10% of all fly brain cells, could also reduce mortality phenotypes.

As shown in Figure 1, we compared survival across a 10-d period in *sgll*^*95*^ flies that carried both the respective gal4 drivers and UAS *hPNPO* constructs to survival in *sgll*^*95*^ flies carrying either the gal4 driver or the UAS *hPNPO* construct alone. In *sgll*^*95*^ flies expressing *hPNPO* in cholinergic neurons (Figure 1A), we observed a marginal increase in survival compared to UAS *hPNPO* only control. (mean survival: 3.4 vs 2.6 d) When *hPNPO* was expressed glutamatergic neurons (Figure 1B), we observed no improvement in mortality (2.0 vs 2.6 d). In contrast, *hPNPO* expression in GABAergic neurons (Figure 1C) substantially improved mortality. Over the 10-day observation period, > 50% of UAS *hPNPO / GAD* gal4 ; *sgll*^*95*^ flies survived, while all control flies died within 5 days (p < 0.0001). Expression of *hPNPO* in glia also improved mortality, albeit to a lesser degree (Figure 1D). We found UAS *hPNPO / repo* gal4 ; *sgll*^*95*^ flies had a mean survival of 6.4 d (p <0.0001). Indeed, a substantial fraction (∼ 20%) of the glia-driven *hPNPO* flies survived beyond 10 d. Our observations indicate *hPNPO* expression in GABAergic neurons is largely sufficient to reduce mortality phenotypes in *sgll*^*95*^ mutants, while glia expression also is beneficial. In contrast, *hPNPO* expression in cholinergic neurons and glutamatergic neurons did not materially affect these phenotypes.

**Figure 1.**
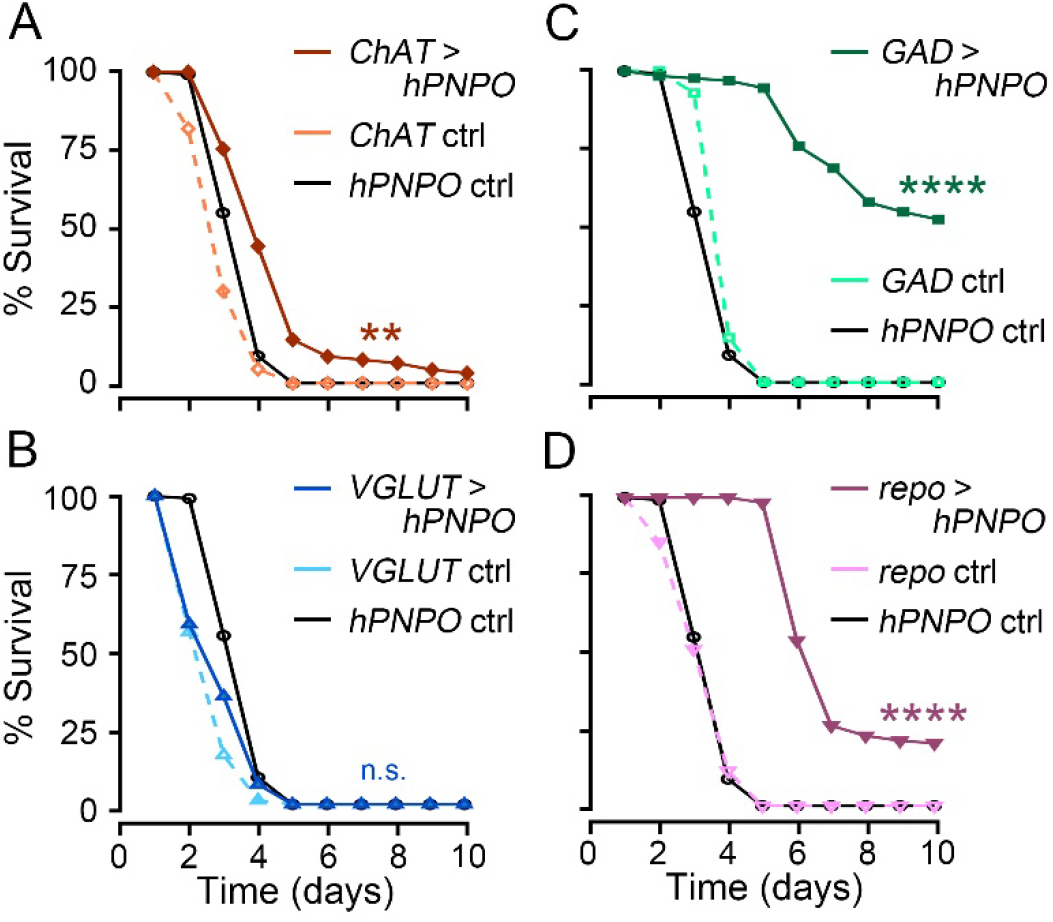
Cell-type specific *hPNPO* expression modulates survival of *sgll*^*95*^ mutants on VB6-free media. Survival curves of *sgll*^*95*^ flies with UAS *hPNPO* driven by selected gal4 lines (i.e. gal4 > *hPNPO*). (A) Cholinergic neurons: UAS *hPNPO / ChAT* gal4 ; *sgll*^*95*^; (B) glutamatergic neurons: UAS *hPNPO / VGLUT* gal4 ; *sgll*^*95*^; (C) GABAergic neurons: UAS *hPNPO / GAD* gal4 ; *sgll*^*95*^; and (D) glia: UAS *hPNPO / repo* gal4 ; *sgll*^*95*^. In each panel, survival curves for the corresponding gal4 driver control line (gal4 / + ; *sgll*^*95*^, dashed line) and *hPNPO* control line (UAS *hPNPO / +* ; *sgll*^*95*^, black line) are plotted. All flies were reared on VB6-free media. N = 75 – 132 flies. ** p < 0.01, **** p < 0.0001 log-rank test vs *hPNPO* control, Bonferroni-corrected.

### Electrophysiological analysis of spontaneous seizures *sgll* mutants expressing *hPNPO*

We sought to determine if spontaneous seizures that manifest in *sgll*^*95*^ flies reared on a VB6-deprived diet would also be affected by *hPNPO* expression. Utilizing a tethered fly preparation (Figure 2A, inset, see Iyengar and Wu 2014[18]), we recorded spiking activity in the dorsal longitudinal flight muscles (DLMs). These large muscles serve as an accessible readout of the spontaneous high-frequency spike bursts synchronized across the left and right motor units that correspond to seizure activity in *sgll* mutants [14, 15].

**Figure 2.**
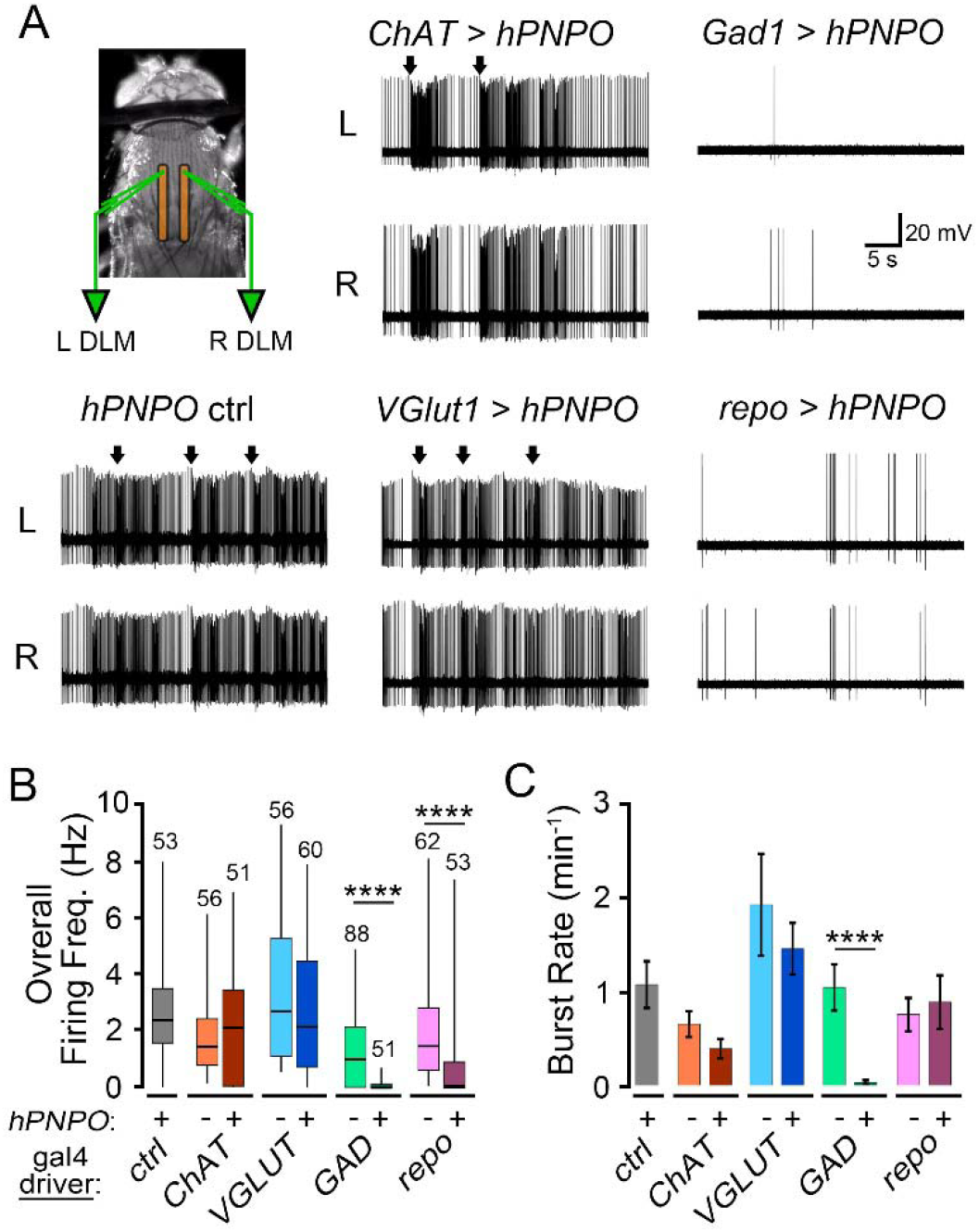
Modulation of spontaneous seizure-associated spike bursts in *sgll*^*95*^ fly mutants via *hPNPO* expression (A) Illustration of tethered fly recording configuration (inset) and representative traces of left (L) and right (R) DLM firing from UAS *hPNPO/+* ; *sgll*^*95*^ control flies (*hPNPO* ctrl) and *hPNPO* driven by respective gal4 drivers on a *sgll*^*95*^ background (i.e. gal4 > *hPNPO*). All flies were reared on VB6-free media. Arrows above traces indicate bilaterally synchronized spike bursts. Note reduced DLM spiking in *GAD* and *repo* driven *hPNPO* expression. (B) Box plots of overall firing frequency (boxes: 25^th^, 50^th^ and 75^th^ %tiles, whiskers: 5^th^ and 95^th^ %tile). (C) Bar graph of average burst rate in the respective genotypes (error bars indicate S.E.M). Sample sizes indicate the number of recordings. *** p < 0.001, **** p < 0.0001, Kruskal-Wallis ANOVA, rank-sum *post hoc* test versus corresponding +/gal4 ; *sgll*^*95*^ control line.

Consistent with our prior reports of *sgll*^*95*^ flies, when reared on VB6-free diets, *sgll*^*95*^ flies carrying either the UAS *hPNPO* or a gal4 construct alone displayed clear spontaneous spike activity, including the characteristic burst phenotype (Figure 1A, UAS *hPNPO / +* ; *sgll*^*95*^ shown). In these control groups, the mean overall firing frequency were between 1.4 – 3.5 Hz (Figure 2B), and the overall burst rate was 0.66 – 1.9 min^-1^ (Figure 2C). When *hPNPO* was expressed in cholinergic or glutamatergic neurons, the DLM spike phenotypes were largely unaltered, with minimal changes in either the overall firing frequency or rate of spike bursts. In contrast, *sgll*^*95*^ flies with *hPNPO* expressed in GABAergic neurons (via *GAD* gal4) displayed markedly improved phenotypes. In these flies, the occasional spontaneous DLM spikes observed corresponded with wing grooming events (Figure 2A). Indeed, the overall DLM firing rate was substantially lower (mean: 0.13 Hz) than either control line, and spike bursts were never observed (Figure 2B-C). Glial *hPNPO* expression (via *repo* gal4) also improved spiking phenotypes in *sgll*^*95*^ flies, with spontaneous spiking (mean: 0.99 Hz) largely reduced. However, in this group, the rescue effect was incomplete, as the spike bursting was not significantly altered compared to gal4 controls (Figure 2C). Overall, our tethered fly results largely mirror our mortality findings, with *hPNPO* expression in GABAergic neurons and glia (to a lesser degree) attenuating seizure phenotypes.

### Modulation of *sgll* mutant mortality via pharmacological manipulation of GABAergic transmission

Based on our findings that *hPNPO* expression in GABAergic neurons markedly improved survival and seizure-related phenotypes in *sgll*^*95*^ mutants, we asked if pharmacological modulation of GABA signaling could also improve mortality in *sgll*^*95*^ flies. We measured the survival of *sgll*^*95*^ flies on VB6-free media containing GABA or one of three GABA receptor modulators: Gaboxadol, a GABA_A_ receptor agonist and GABA structural analog previously studied as a sleep-promoting drug in flies [21]; Diazepam, a benzodiazepine and GABA_A_ positive allosteric modulator commonly used to treat status epilepticus, that can suppress seizures in flies [22]; and SKF-97541, a GABA_B_ receptor agonist, also known to affect sleep architecture [23]. As shown in Figure 3, none of these drugs recapitulated improvements when *hPNPO* was expressed in GABAergic neurons. Indeed, only SKF-97541 feeding resulted in a significant, but modest survival extension (3.7 vs 2.7 d when fed 1.0 µg/ml), while feeding GABA or the GABA_A_ modulators did not significantly reduce mortality (Figure 3). At the respective concentrations, control *w*^*1118*^ flies did not show any mortality over the observation period (i.e. 100% survival). These drug-feeding experimental results indicate a potential contribution for loss of GABA_B_ signaling in *sgll* mutant phenotypes.

**Figure 3.**
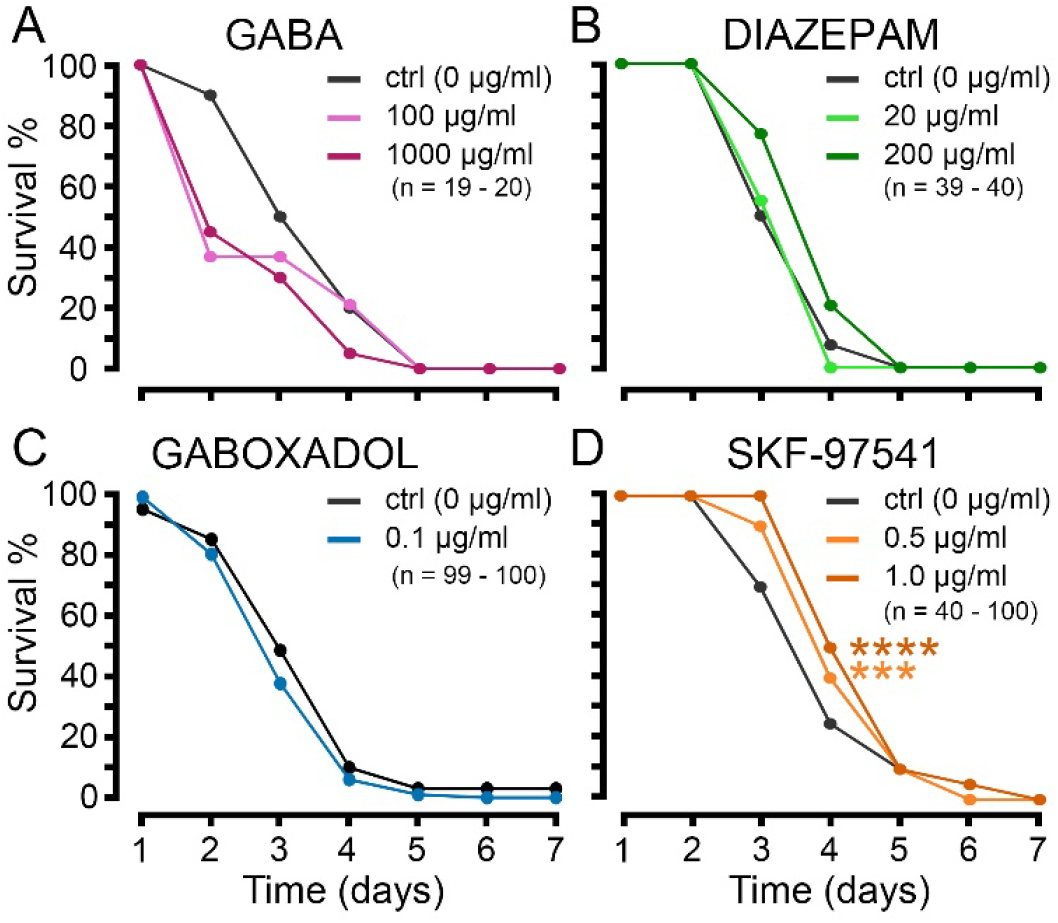
Effect of feeding GABAergic modulators on mortality in VB6-deficient *sgll*^*95*^ flies. Following eclosion, *sgll*^*95*^ flies were reared on VB6-free media containing the following compounds: (A) GABA, (B) diazepam, (C) gaboxadol, (D) SKF-97541 at the indicated concentrations. Survival over a 7-d period is plotted as in Figure 1. Sample sizes (number of flies) as indicated.*** p < 0.001, **** p < 0.0001; log-rank test vs. ctrl, Bonferroni-corrected.

## Discussion

Mutations in *PNPO* are identified as a relatively common cause for epilepsy syndromes in humans [24]. Seizures in patients carrying PNPO mutations are largely resistant to traditional anti-seizure medications. In most individuals, VB6 (PN and/or PLP) dietary therapies are usually effective, although seizure control is sometimes incomplete [10, 25, 26]. We have previously shown seizure phenotypes, including convulsions and spontaneous spike discharges, manifest in flies carrying loss-of-function mutations *sgll* (the sole *PNPO* gene) when they are reared on a VB6 vitamer-deprived diet [14]. Transgenic expression of *hPNPO* in *sgll* mutants reversed these phenotypes—indicating human PNPO can functionally replace the fly enzyme. Furthermore, in flies where the *sgll* locus was replaced with epilepsy-associated *hPNPO* alleles, range of seizure phenotypes emerged which was reminiscent of variability across patient population [15]. Thus, PNPO deficiency-linked epilepsy can be modeled effectively in *Drosophila*.

This study compared how transgenic expression of *hPNPO* in different cell-types of the nervous system affected seizure expression in *sgll* mutants. We found that *hPNPO* expression in GABAergic neurons, and to a lesser extent glia improved both survival (Figure 1) and electrophysiological (Figure 2) phenotypes. In contrast, *hPNPO* expression in excitatory cholinergic neurons (accounting for most synapses in the fly brain [27]) or glutamatergic neurons had minor phenotypic effects. Given the pivotal role of PLP in GABA synthesis, it is likely reduced inhibitory GABAergic transmission is a major contributing factor to the emergence of seizures in the context of PNPO deficiency. Indeed, zebrafish *pnpo*^*-/-*^ larvae have substantially reduced GABA levels compared to control [13]. Seizure-related spike discharges in fly *sgll* mutants are reminiscent of discharges associated with GABA_A_ receptor blockade via picrotoxin [20], and these mutants have a lower threshold for picrotoxin response [15]. Our findings indicate direct cell-autonomous PLP synthesis in GABAergic neurons can be sufficient to improve outcomes in *PNPO* mutants, further highlighting the specific role of GABAergic circuitry in underlying PNPO-deficiency related epilepsy.

While PNPO activity in GABAergic neurons may produce enough PLP to attenuate seizure expression in *sgll* mutants, it is notable that neither pan-neuronal nor GABAergic neuron-specific *sgll* RNAi knock-down lead to clear seizure phenotypes (Chi et al., 2019; data not shown). In GABAergic neurons with deficient PNPO activity, a non-cell autonomous pathway consisting of uptake of circulating PLP or PL vitamers may provide a sufficient source. (Notably, PL to PLP conversion is independent of PNPO). We observed moderate phenotypic improvements in *sgll* mutants with glial PNPO expression, and minimal changes when PNPO was expressed in cholinergic or glutamatergic neurons (Figure 1 & 2). As high levels of free B6 vitamers can be toxic [28], circulating PLP levels are tightly regulated by sequestration to PLP binding proteins, and dephosphorylation by non-specific phospshatases [5]. The close physical proximity and extensive inter-cellular transport pathways between glia and GABAergic neurons could overcome these regulatory processes and shuttle B6 vitamers in a way that surround neurons cannot. Thus, PNPO function in glia could contribute to locally circulating B6 vitamer levels to support GABAergic neuron function. However, the glial contribution is likely limited, as direct PLP feeding appears more effective at seizure suppression (compare Figure 1D vs Figure 4 in Chi et al., 2014).

The improved survival and attenuated seizure expression in *sgll* mutants following hPNPO expression in GABAergic neurons prompted us to examine how pharmacological modulation of GABA signaling impacted survival phenotypes. We adopted a drug feeding approach, using compounds with demonstrated effects on ionotropic GABA_A_ or metabotropic GABA_B_ receptor function in *Drosophila*. We found neither GABA_A_ receptor agonists GABA and gaboxadol nor the GABA_A_ positive allosteric modulator diazepam increased survival (Figure 3A-C). Our observations appear consistent with clinical reports of benzodiazepines (including diazepam) in patients carrying *PNPO* mutations, where these drugs suppress acute seizures but are not effective in long-term seizure control [29, 30]. Feeding the GABA_B_ receptor agonist SKF-97541 had the greatest effect amongst the drugs tested, improving survival by ∼ 1 d (Figure 3D). In *Drosophila*, GABA_B_ receptors modulate sleep architecture and sensitivity to alcohol, much like their role in vertebrates [31-33]. Disruption of GABA_B_ signaling is an important factor in a variety of epilepsy syndromes [34, 35]. Our observations may suggest reduced activation of GABA_B_ receptors may contribute to seizure phenotypes arising from *PNPO* mutations.

In summary, our studies indicate seizure phenotypes in *sgll* mutant induced by VB6 deprivation can be ameliorated by PNPO expression in GABAergic neurons and in glia (to a lesser degree). The effects of targeted PNPO expression are stronger than direct pharmacological manipulations of GABAergic signaling. Our work also demonstrates the utility of fly *sgll* mutants as a model system for establishing mechanistic links between genotype, dietary factors and phenotype in the context of VB6 vitamer and/or PNPO deficiency.

## Acknowledgments

We thank Wanhao Chi for input during early stages of the project; Reid Schuback for supporting electrophysiology experiments; and members of the Zhuang and Iyengar labs for their technical assistance, helpful comments and encouragement. This project was supported by NIH Grants: NS 111122 (to XZ), and NS 134960 (to AI) as well as U. Alabama start-up funds (to AI)

## Notes

### Competing Interest Statement

The authors have declared no competing interest.

## References

1. Fitzpatrick, Teresa B., et al., Two independent routes of de novo vitamin B6 biosynthesis: not that different after all. Biochemical Journal, 2007. 407(1): p. 1–13.

2. Ueland, P.M., et al., Direct and Functional Biomarkers of Vitamin B6 Status. Annual Review of Nutrition, 2015. 35(Volume 35, 2015): p. 33–70.

3. Parra, M., S. Stahl, and H. Hellmann, Vitamin B6 and Its Role in Cell Metabolism and Physiology. Cells, 2018. 7(7): p. 84.

4. Rivero, M., N. Novo, and M. Medina, Pyridoxal 5′-Phosphate Biosynthesis by Pyridox(am)-ine 5′-Phosphate Oxidase: Species-Specific Features. International Journal of Molecular Sciences, 2024. 25(6): p. 3174.

5. di Salvo, M.L., R. Contestabile, and M.K. Safo, Vitamin B6 salvage enzymes: Mechanism, structure and regulation. Biochimica et Biophysica Acta (BBA) - Proteins and Proteomics, 2011. 1814(11): p. 1597–1608.

6. Percudani, R. and A. Peracchi, A genomic overview of pyridoxalJphosphateJdependent enzymes. EMBO reports, 2003. 4(9): p. 850–854.

7. Plecko, B. and P. Mills, PNPO Deficiency. 1993: University of Washington, Seattle, Seattle (WA).

8. Plecko, B., et al., Pyridoxine responsiveness in novel mutations of the PNPO gene. Neurology, 2014. 82(16): p. 1425–1433.

9. Guerriero, R.M., et al., Systemic Manifestations in Pyridox(am)ine 5′-Phosphate Oxidase Deficiency. Pediatric Neurology, 2017. 76: p. 47–53.

10. Alghamdi, M., et al., Phenotypic and molecular spectrum of pyridoxamine-5′-phosphate oxidase deficiency: A scoping review of 87 cases of pyridoxamine-5′-phosphate oxidase deficiency. Clinical Genetics, 2021. 99(1): p. 99–110.

11. Mastrangelo, M., et al., Epilepsy Phenotypes of Vitamin B6-Dependent Diseases: An Updated Systematic Review. Children, 2023. 10(3): p. 553.

12. Chen, P.-Y., et al., Pyridoxamine Supplementation Effectively Reverses the Abnormal Phenotypes of Zebrafish Larvae With PNPO Deficiency. Frontiers in Pharmacology, 2019. Volume 10 - 2019.

13. Ciapaite, J., et al., Pyridox(am)ine 5′-phosphate oxidase (PNPO) deficiency in zebrafish results in fatal seizures and metabolic aberrations. Biochimica et Biophysica Acta (BBA) - Molecular Basis of Disease, 2020. 1866(3): p. 165607.

14. Chi, W., et al., Pyridox (am) ine 5’-phosphate oxidase deficiency induces seizures in Drosophila melanogaster. Human Molecular Genetics, 2019. 28(18): p. 3126–3136.

15. Chi, W., et al., Drosophila carrying epilepsy-associated variants in the vitamin B6 metabolism gene PNPO display allele- and diet-dependent phenotypes. Proceedings of the National Academy of Sciences, 2022. 119(9): p. e2115524119.

16. Brand, A.H. and N. Perrimon, Targeted gene expression as a means of altering cell fates and generating dominant phenotypes. Development, 1993. 118(2): p. 401–415.

17. Chi, W., et al., A Nutritional Conditional Lethal Mutant Due to Pyridoxine 5′-Phosphate Oxidase Deficiency in Drosophila melanogaster. G3 Genes|Genomes|Genetics, 2014. 4(6): p. 1147–1154.

18. Iyengar, A. and C.-F. Wu, Flight and Seizure Motor Patterns in Drosophila Mutants: Simultaneous Acoustic and Electrophysiological Recordings of Wing Beats and Flight Muscle Activity. Journal of Neurogenetics, 2014. 28(3-4): p. 316–328.

19. Miller, A., The Internal Anatomy and Histology of the Imago of Drosophila melanogaster, in Biology of Drosophila, M. Demerec, Editor. 1950, Cold Spring Harbor Press. p. 420–534.

20. Lee, J., A. Iyengar, and C.-F. Wu, Distinctions among electroconvulsion-and proconvulsant-induced seizure discharges and native motor patterns during flight and grooming: quantitative spike pattern analysis in Drosophila flight muscles. Journal of neurogenetics, 2019. 33(2): p. 125–142.

21. Dissel, S., et al., Sleep Restores Behavioral Plasticity to <em>Drosophila</em> Mutants. Current Biology, 2015. 25(10): p. 1270–1281.

22. Stilwell, G.E., et al., Development of a Drosophila seizure model for in vivo high-throughput drug screening. Eur J Neurosci, 2006. 24(8): p. 2211–22.

23. Kim, M., et al., Rogdi Defines GABAergic Control of a Wake-promoting Dopaminergic Pathway to Sustain Sleep in Drosophila. Scientific Reports, 2017. 7(1): p. 11368.

24. Abou-Khalil, B., et al., Genome-wide mega-analysis identifies 16 loci and highlights diverse biological mechanisms in the common epilepsies. Nature Communications, 2018. 9(1): p. 5269.

25. Jiao, X., et al., Analysis for variable manifestations and molecular characteristics of pyridox(am)ine-5′-phosphate oxidase (PNPO) deficiency. Human Molecular Genetics, 2022. 32(11): p. 1765–1771.

26. Mills, P.B., et al., Epilepsy due to PNPO mutations: genotype, environment and treatment affect presentation and outcome. Brain, 2014. 137(5): p. 1350–1360.

27. Lin, A., et al., Network statistics of the whole-brain connectome of Drosophila. Nature, 2024. 634(8032): p. 153–165.

28. Hadtstein, F. and M. Vrolijk, Vitamin B-6-Induced Neuropathy: Exploring the Mechanisms of Pyridoxine Toxicity. Advances in Nutrition, 2021. 12(5): p. 1911–1929.

29. Jensen, K.V., et al., Diagnostic pitfalls in vitamin B6-dependent epilepsy caused by mutations in the PLPBP gene. JIMD Reports, 2019. 50(1): p. 1–8.

30. Levtova, A., et al., Normal Cerebrospinal Fluid Pyridoxal 5′-Phosphate Level in a PNPO-Deficient Patient with Neonatal-Onset Epileptic Encephalopathy, in JIMD Reports, Volume 22, J. Zschocke, et al., Editors. 2015, Springer Berlin Heidelberg: Berlin, Heidelberg. p. 67–75.

31. Gmeiner, F., et al., GABAB receptors play an essential role in maintaining sleep during the second half of the night in Drosophila melanogaster. Journal of Experimental Biology, 2013. 216(20): p. 3837–3843.

32. Dzitoyeva, S., N. Dimitrijevic, and H. Manev, γ-Aminobutyric acid B receptor 1 mediates behavior-impairing actions of alcohol in Drosophila: Adult RNA interference and pharmacological evidence. Proceedings of the National Academy of Sciences, 2003. 100(9): p. 5485–5490.

33. Ranson, D.C., et al., Pharmacological targeting of the GABAB receptor alters Drosophila’s behavioural responses to alcohol. Addiction Biology, 2020. 25(2): p. e12725.

34. Joshi, K., M.A. Cortez, and O.C. Snead, Targeting the GABAB Receptor for the Treatment of Epilepsy, in GABAB Receptor, G. Colombo, Editor. 2016, Springer International Publishing: Cham. p. 175–195.

35. Han, H.A., M.A. Cortez, and O.C. Snead, III, GABA(B) Receptor and Absence Epilepsy, in Jasper’s Basic Mechanisms of the Epilepsies, J.L. Noebels, et al., Editors. 2012, National Center for Biotechnology Information (US)

